# Lateralized dopaminergic neuron activity encodes goal-directed turning

**DOI:** 10.64898/2026.09.23.753763

**Authors:** Ashwin Miriyala, Scott Waddell

## Abstract

Animals navigate through dynamic environments in search of resources. Dopaminergic systems are known to control movement and assign motivational value to cues, actions and outcomes, but how they guide ongoing locomotion towards a goal remains unclear. Here, we demonstrate that Drosophila mushroom body innervating dopaminergic neurons encode steering signals in world-centred (allocentric) and body-centred (egocentric) coordinates during goal-directed navigation. Thirsty flies track water vapour through upwind directed turning and locomotion, that persists for several seconds after the vapour disappears. We identified a population of rewarding dopaminergic neurons to be specifically required for maintaining directed tracking following vapour offset. Recordings of these dopaminergic neurons during behaviour revealed delayed bilateral activity that followed upwind-directed turning in allocentric coordinates. Moreover, lateralized ipsilateral activity in a subset of these dopaminergic neurons encode whether the fly turns left or right in egocentric coordinates. Finally, these dopaminergic neurons also exhibited increased ipsilateral activity when turning in the direction where water vapour was previously encountered. Therefore, activity of mushroom body dopaminergic neurons represents steering and direction-specific actions that together maintain goal-directed tracking in turbulent environments.

## Introduction

Animals display remarkable feats of navigation as they search for resources like food and water. Ants integrate paths between visual landmarks on a wandering outward journey to generate a homing vector to directly return to their nest ^1^ . Albatross and shearwaters use olfactory cues in a visually featureless expanse of sea to locate food ^2,3^. Studying such navigational strategies can reveal how neuronal circuits perform fundamental computations.

Using airborne smells to search for resources poses many challenges, such as extracting meaning from transient encounters with a turbulent plume, to inform ongoing behavioural adjustments ^4–7^. When navigating dynamic olfactory environments, wind provides information of source direction while odours can provide value signals that inform the fly that it is tracking something meaningful. An unprecedented view of the neuronal computations involved during navigation have come from studies of the locust ^8^, butterfly ^9^ and *Drosophila melanogaster* central complex (CX). The insect CX provides a compass system that computes differences between allocentric (world-centered) goal direction vectors and egocentric (self-centered) heading vectors to drive steering ^10–17^. Importantly, neuronal networks also need to track action and appropriate outcome to determine whether progress is being made towards a goal. Whether and where these computations exist to complement the function of regions like the CX remains untested.

The *Drosophila* mushroom bodies are a bilaterally symmetric network in which state-dependent innate and learned values can be assigned odours. Dense output connectivity to the CX compass system suggests these values are available to aid resource seeking, but current evidence is limited. Odors are uniquely represented as activity in sparse subsets of the larger population of intrinsic mushroom body neurons, Kenyon Cells (KCs), in each hemisphere of the fly’s brain. A specific stream of α′β′ KCs is dedicated to the thermo-hygrosensory modality, emphasizing the importance of water vapour as a reliable sign of nearby water. Axons of these and other KCs in the mushroom body lobes are tiled by different types of dopaminergic neurons (DANs), which can reinforce memories by directing plasticity of KC synapses onto downstream mushroom body output neurons (MBONs), and also directly modulate MBONs.

Critically, some mushroom body DAN types show responses to movement ^18^ in addition to external reinforcers, making it possible that the mushroom bodies also evaluate locomotor actions to guide navigation towards a goal. Here we directly tested whether mushroom body DANs can provide action-outcome signals by monitoring their activity in thirsty flies searching for water.

## Results

### Thirsty flies navigate upwind towards water vapour

To study goal directed navigation, we pin-tethered individual water-deprived flies and allowed them to walk on an air-cushioned trackball, while delivering water vapour in open-loop into a constant airstream (Figure 1a, Supplementary Figure 1a-b). Direction of airflow was yoked to the fly’s heading direction, within -75 to +75 degree limits centered on the fly’s forward axis (Figure 1b). This allowed us to measure goal-directed approach as the fly navigated within an air stream.

**Figure 1.**
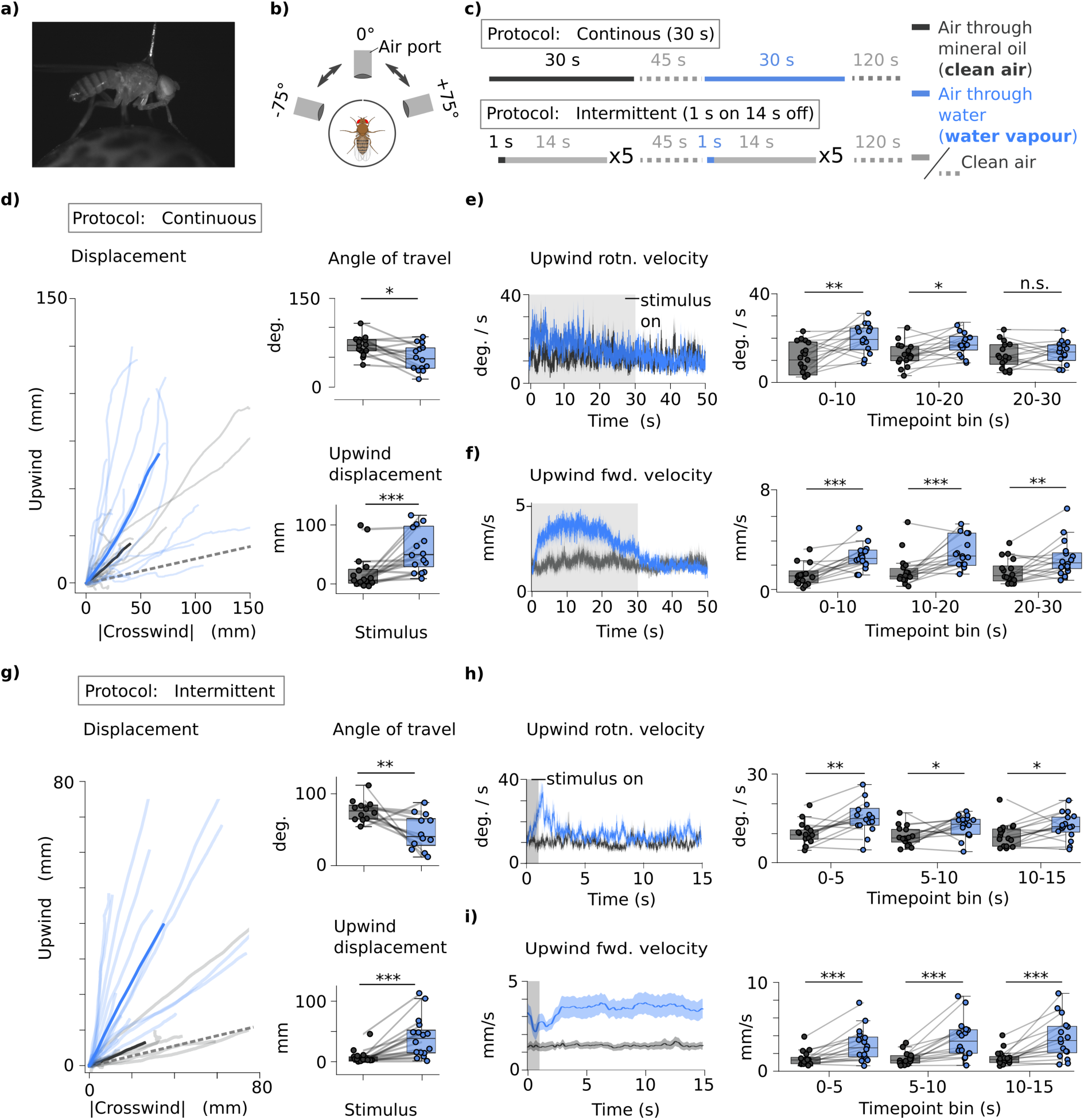
Thirsty flies navigate upwind towards water vapour. **a,** Side view of a pin-tethered fly walking on a trackball. **b,** Cartoon of closed-loop stimulus delivery. **c,** Description of the continuous (30 s) and intermittent (1 s on 14 s off) protocols. Air bubbled through either water (vapour, solid blue lines) or mineral oil (clean air, solid grey lines) replaced constant airflow (dotted lines). **d, left:** Walking trajectories during continuous protocol, plotted as upwind (positive y-axis) and absolute crosswind (x-axis) position over the 30 clean air (gray) and vapour (blue) exposures. Dashed grey line represents servo limit at 75 °. Transparent lines are average trajectory per fly while solid lines represent trajectory averaged across flies. **right:** In continuous (30 s) protocol, flies travel upwind at a significantly smaller angle of travel (top; n=14 flies) and increased displacement (bottom; n=15 flies) when vapour is on as compared to in clean air. **e,** Upwind rotational (rotn.) velocity during vapour exposure (grey region) reveals upwind turning during the first 20 s of water vapour exposure (n=15 flies). **f,** Upwind forward (fwd.) velocity is elevated throughout the 30 s vapour exposure and returns to baseline with vapour offset. **g,** Trajectory during intermittent protocol (1 s on 14 s off; averaged across five vapour exposures and two protocol repeats, then across flies). Over the 15 s duration, flies travel upwind at a reduced angle of travel (n=12 flies) and increased displacement (n=16 flies) as compared to in clean air. **h-i,** Upwind rotational velocity (h) and upwind forward velocity (i) are elevated during and after vapour exposure (n=16 flies). All comparisons between clean air and water vapour groups were compared using a two-sided paired Wilcoxon signed-rank test. All boxplots show median and interquartile range (IQR), and whiskers represent data within 1.5 * IQR. Exact statistical values and comparisons are presented in Supplementary Information.

Since insects track olfactory plumes by integrating information across intermittent exposures, that can be of variable duration ^5–7,19^, we delivered water vapour with two distinct temporally structured protocols (Figure 1c). In the continuous protocol, flies experienced a continuous 30 s stimulus exposure, which was repeated 4 times per fly with 2 min rest intervals. In the intermittent protocol, flies experienced five intermittent 1 s stimulus exposures with 14 s inter stimulus intervals of ‘clean air’, which was repeated 2 times per fly with a 2 min rest interval. Vapour stimulus was generated by bubbling air through water, while air passed through mineral oil served as the ‘clean air’ stimulus. Statistical comparisons of vapour-evoked behaviour were made relative to behaviour measured during the clean air stimulus within each protocol. Behaviour from a protocol repeat was only analyzed if the fly walked an average of 3 mm/s over the protocol duration.

Behaviour was quantified over the stimulus period by calculating the upwind angle of travel (angle of the upwind x |crosswind| running trajectory relative to the vertical y-axis, such that a 0° angle indicates the fly is running upwind into the airstream) and the upwind displacement (the final position in mm along the vertical y-axis). Behaviour measured using the continuous protocol (averaged across all 4 protocol repeats; Figure 1d) showed that in clean air, flies travel upwind at a median angle of 71 degrees (IQR=19) with a displacement of 7 mm (IQR=23). Whereas, when water vapour was turned on, flies showed clear signs of stimulus tracking. Upwind heading angle was significantly reduced (48, IQR=32) while upwind displacement increased (51 mm, IQR= 69) relative to clean air. Therefore, pin-tethered flies on a trackball exhibit robust upwind approach in the presence of vapour, consistent with behaviour of freely moving flies walking in an odour stream ^14^.

Approach to water vapour is thirst state dependent, with sated or hungry flies avoiding water vapour in a two-choice assay ^20^ (Supplementary Figure 1c). We therefore verified that vapour approach of individual flies on the trackball also showed state-dependence. Exposing hungry flies to water vapour in both the continuous and intermittent protocols on the trackball did not evoke an upwind response distinguishable from their behaviour in clean air (Supplementary Figure 1d). We could not compare behaviour to that of sated flies because sated flies generally show lower levels of locomotion, likely due to lack of motivation ^21^. Upwind approach to water vapour of tethered flies on the trackball is therefore thirst state-specific.

During olfactory navigation, animals continuously sample their environment by sniffing, or casting from side to side ^4^. We therefore extracted rotational information over time in the continuous protocol (Figure 1e) to determine how upwind heading was affected by water vapour. These analyses revealed that flies on average make spontaneous upwind rotations of ∼12 degrees every second in clean air. In contrast, when exposed to water vapour, flies turn upwind more frequently at 20 °/s for 10 s, then gradually revert to ∼12 °/s after 20 s. In addition, quantifying upwind velocity showed flies move faster at ∼3 mm/s while exposed to water vapour, and quickly return to baseline velocity of ∼ 1 mm/s after vapour offset (Figure 1f). The behavioural response to water vapour did not significantly vary across the four continuous protocol repeats, showing that habituation does not occur with this stimulus protocol (Supplementary Figure 1e). Furthermore, behavioural metrics prior to stimulus onset were not different between clean air and water vapour groups (Supplementary Figure 1f). Together these data show that thirsty flies exhibit spontaneous upwind turning and locomotion in clean air and that continuous exposure to water vapour increases velocity and the frequency of upwind turning for up to 30 s.

Robust increases in upwind approach were also observed when flies were presented with the intermittent (1 s on 14 s off) protocol (Figure 1g). Over 15 s of clean air (averaged across the five exposures and two protocol repeats), flies travelled upwind at 65 degrees (IQR=24) with a displacement of 13 mm (IQR=37). Whereas, over the 15 s of an intermittent trial, flies travelled upwind at a significantly reduced angle (44 degrees, IQR=29) and greater displacement (188 mm, IQR=225).

Although behaviour during vapour was similar between continuous and intermittent protocols, behaviour was radically different after vapour offset (Supplementary Figure 1g). In the continuous protocol, averaged behaviour across flies reveals upwind and rotation returned to baseline upon vapour offset (Figure 1e). However, averaged behaviour in the intermittent protocol, upwind approach initiated during the 1 s vapour exposure continued over the 14 s following vapour offset. Persistent upwind pursuit in the intermittent protocol was also observed as an elevated upwind rotation of 10-15 °/s on average (Figure 1h) alongside an elevated upwind velocity of around 2-4 mm/s on average (Figure 1i) over the entire 15 s (1 s on 14 s off) duration. Additionally, the magnitude of upwind approach increased between the first and second protocol repeats (but not across the five exposures within an individual protocol repeat; Supplementary Figures 1h-j). This increased upwind locomotion between repeats is consistent with prior reports of behaviour in freely moving flies experiencing brief odour exposures ^6,7,22^. Behaviour prior to any stimulus exposure was not different between clean air and water vapour groups (Supplementary Figure 1i). These intermittent protocol data reveal that flies track reward predictive vapour and suggest that they retain a memory of brief, intermittent encounters to maintain directed pursuit when they lose track of vapour.

Taken together, these data reveal that thirsty flies exhibit robust upwind turning and locomotion when they detect water vapour and that if the experience is fleeting, they continue tracking in the same direction. Importantly, using the two temporal structures of the continuous and intermittent protocols permits study of approach when vapour is present and the maintenance of approach when the vapour disappears. We next tested whether the mushroom bodies instruct upwind approach to water vapour.

### A subset of reward DANs are required to maintain approach following vapour offset

Prior studies have suggested that different mushrooom body DAN types reinforce water learning from those controlling innate water seeking ^23,24^. We therefore used the R48B04-GAL4 driver that labels some learning and seeking DAN types (γ4, γ5 and β′2) to test whether these neurons mediate upwind approach to water vapour in individual thirsty flies. We blocked output from R48B04 DANs using expression of the dominant temperature-sensitive UAS-*Shibire*^ts^^1^ (*Shi*^ts1^) transgene ^25^. *Shi*^ts1^ blocks membrane recycling and thus synaptic vesicle release at the restrictive temperature > 29 °C and this blockade is reversible by returning flies to 25 °C. A wire heater set to 32 °C was immersed in a droplet of saline placed on the head of a head-fixed fly, and behaviour was measured while the fly navigated the continuous and intermittent water vapour exposure protocols on a trackball. In these intervention experiments we tested behaviour in a 15 s continuous protocol because upwind rotation and velocity of wild-type flies were strongest in the first half of the continuous 30 s exposure (Figure 1e).

Unexpectedly, blocking the R48B04 DANs did not alter upwind approach during the continuous exposure protocol. Upwind angle of travel and displacement was similar to that of genetic control flies (Figure 2a) and all flies showed clear signs of vapour tracking relative to their behaviour in a clean air stream (Figure 2b). Instead, vapour tracking was clearly impaired when the DANs were blocked throughout navigation in the intermittent protocol. DAN blocked flies travelled upwind at an increased angle and reduced displacement over the 1 s on 14 s off duration (averaged over all five exposures and across protocol repeats), as compared to their genetic controls (Figure 2c). Behaviour in clean air of all genotypes was similar at restrictive temperature (Supplementary Figure 2a), and upwind tracking of water vapour was similar across genotypes at permissive 23 °C (Supplementary Figure 2b), ruling out temperature induced motor defects.

**Figure 2.**
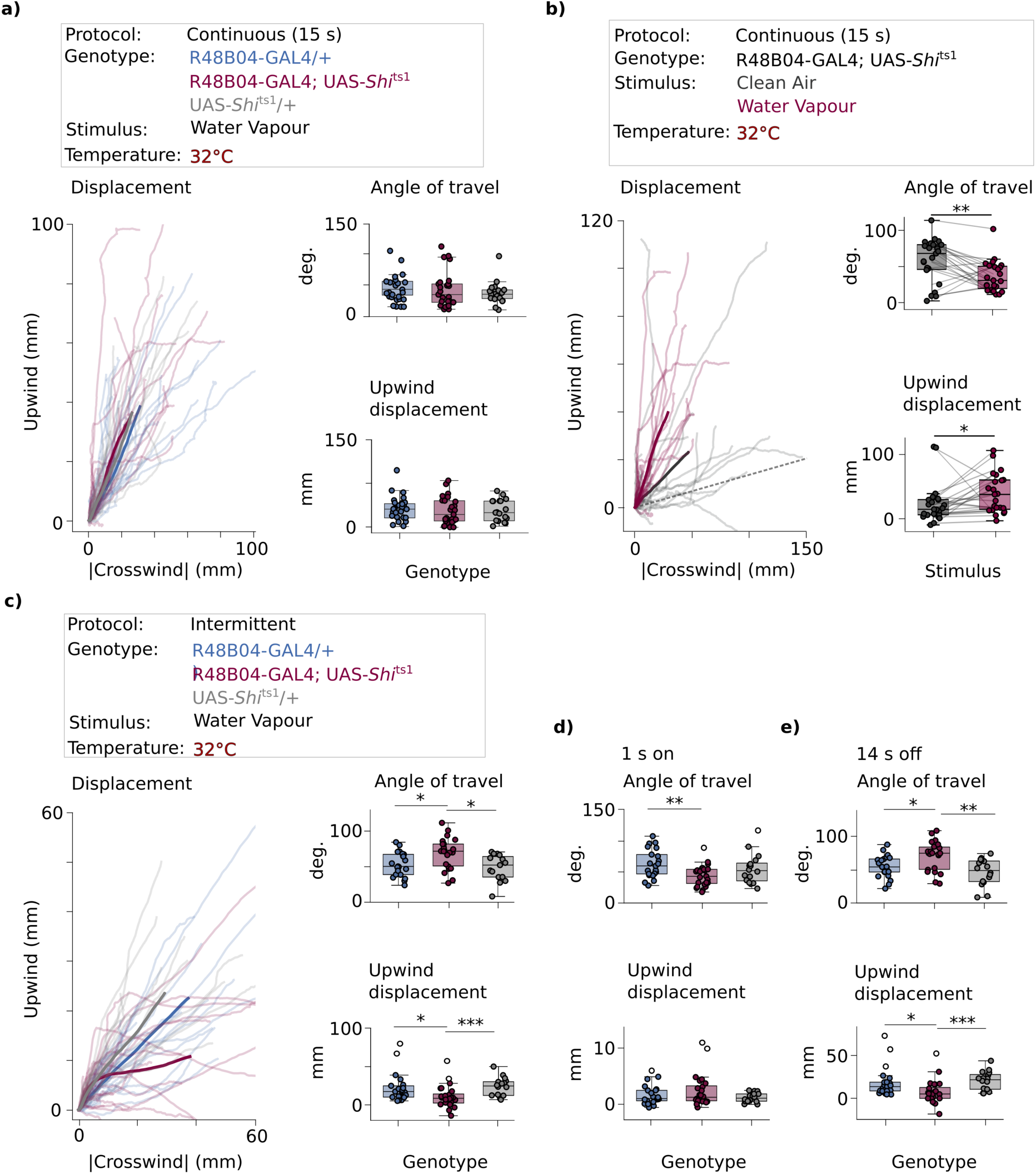
DAN output maintains upwind approach following vapour offset. **a-b,** Blocking R48B04 neurons does not impair upwind approach in the continuous (15 s) protocol when measured a) relative to control genotypes (n=18-25 flies for each group and b) relative to clean air (n=22-23 flies for each paired group; two-sided paired Wilcoxon signed-rank test). **c**) R48B04 neuron block impairs upwind approach in the intermittent protocol, relative to control genotypes (behaviour averaged across 5 exposures and over two protocol repeats; n=19-24 flies per group). **d-e,** Approach is not impaired during 1 s vapour exposure (d) but is impaired following vapour offset (e) as measured relative to control genotypes (n=19-24 flies per group). Hollow dots indicate outliers that are > 1.5 * inter-quartile range. Unless otherwise stated, statistical comparisons between groups were made using a Kruskal-Wallis test followed by Dunn’s multiple comparisons test with Bonferroni correction. Exact statistical values and comparisons are presented in Supplementary Information.

Since wild-type flies initiated approach during the 1 s vapour exposure and maintained approach for at least 14 s afterwards (Supplementary Figure 1e), we tested whether DANs might be differentially required while tracking vapour on and vapour off durations. Surprisingly, analyses of behaviour during these sections of the intermittent protocol revealed the DANs to be dispensable for tracking in the presence of vapour but to be required to maintain tracking when vapour is lost (Figure 2d,e). Following vapour offset, DAN blocked flies travelled at a significantly larger angle and shorter displacement relative to their genetic controls. We replicated this defect in the maintenance of tracking using UAS-*Shibire*^ts1^ driven by 0104-GAL4 (Supplementary figure 2c-e), which expresses in overlapping populations of β′2 DANs, and partially overlapping γ4 DANs to that of R48B04, but largely different subsets of γ5 DANs ^23^. The intersection of these loss-of-function experiments therefore suggests that γ4, γ5 and β′2 DAN activity is dispensable for tracking when the vapour is present, but γ4 and β′2 DANs are required when vapour is lost. These data are consistent with memory of vapour history being required to maintain an appropriate approach trajectory.

### Upwind locomotion is correlated with γ4 and β′2 DAN activity

To further understand how γ4 and β′2 DANs aid vapour tracking, we monitored their activity (with γ5 as control) while measuring behaviour. In these experiments we exclusively used the intermittent protocol, so that we could measure how DANs responded in the presence of vapour and when it is lost, when their output is required.

Individual water-deprived flies that expressed UAS-GCaMP8f; tdTomato in R48B04-GAL4 neurons were head-fixed and their brains were imaged under two-photon illumination while they navigated on the trackball in the intermittent protocol. Signals were simultaneously recorded from both sides of the brain at the level of mushroom body lobe compartments where they can be assigned to the presynaptic fields of individual DAN types (Figure 3a). Co-expression of red fluorescent tdTomato provided reference for correction of brain movement. As before with thirsty pin-tethered flies, flies head-fixed under the microscope showed clear initiation of upwind tracking behaviour during the 1 s vapour exposure and maintained tracking during the 14 s off period (averaged across the 5 exposures and two protocol repeats per fly; Figure 3b). Flies travelled upwind at reduced angle in both 1 s vapour on (median=50, IQR=31) and 14 s vapour off (median=34, IQR=23), relative to their angle of travel in clean air (1 s on: median=73, IQR=19; 14 s off: median=53, IQR=18). While head-fixed flies travelled further upwind during 1 s vapour on relative to clean air (water vapour: median displacement=2 mm, IQR=2 mm; clean air: median=1 mm, IQR=1 mm), there was no difference in their tracking during the 14 s off period, likely because flies are generally more active under 2-photon illumination (median=17, IQR=18 for both clean air and water vapour).

**Figure 3.**
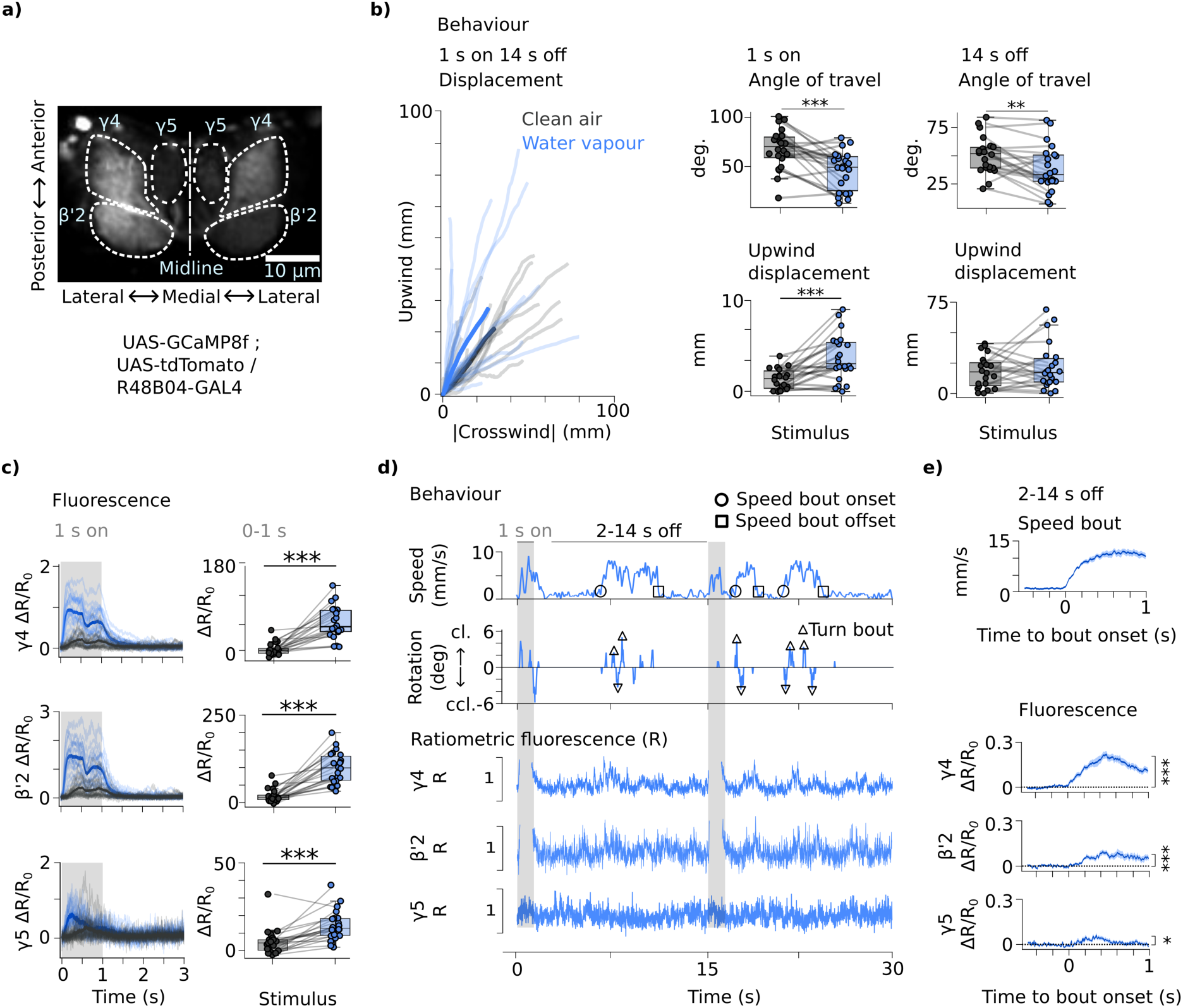
Vapour and locomotion evoke DAN activity. **a, Representative** two-photon image showing regions of interest segmenting signals for DAN types co-expressing GCaMP8f (displayed) and tdTomato. **b) left:** Trajectories of head-fixed, thirsty flies navigating the intermittent protocol under two-photon illumination. **middle and right:** During 1 s vapour exposure (middle column), flies walk upwind at significantly lower angle of travel and increased displacement and in the 14 s off duration (right column) walk at significantly reduced angle of travel (n=22 flies). **c,** Water vapour-triggered γ4, β′2 andγ5 DAN activity (R=ratiometric fluorescence, calculated as GCaMP fluorescence / tdTomato fluorescence). Area under curve measured from the 0-1 s stimulus duration (n=22 flies). **d,** Example speed and rotation (clockwise; cl as positive values and counterclockwise; ccl as negative values) alongside corresponding γ4, β′2 and γ5 DAN fluorescence from one fly over two 1 s water vapour exposures (grey bars) separated by 14 s of clean air. Circles and squares indicate onset and offset of speed bouts, respectively. Triangles represent detected turn bouts. Fluorescence during 1 s water vapour exposures is clipped on the y-axis. **e, top:** Average trace of speed bouts extracted from the 2-14 s following vapour offset and **bottom:** associated γ4, β′2 and γ5 DAN fluorescence (n=16 flies). Significance from zero measured from area under curve in 0-1 s. Traces plotted over time as mean ± SEM. For b-c, transparent traces are responses for individual flies and solid trace is average across flies. Statistical comparisons made using two-sided paired Wilcoxon signed-rank test. Exact statistical values and comparisons are presented in Supplementary Information.

Although previous studies reported β′2, γ4 and γ5 DANs to respond to water drinking in thirsty flies ^23,26^, their response to vapour has not been established. We found that 1 s water vapour exposure evoked a significant bilateral increase in activity across all three DAN types that returned to baseline within 2 s following vapour offset (Figure 3c, Supplementary Fig 3a). The apparently bi-phasic appearance of DAN activity during vapour exposure results from minor fluctuations in air flow that accompanies the toggling of air valves, as evidenced by their presence in recordings during clean air that involve the same valve operations (Figure 3c; Supplementary Figure 1a; also see methods). Importantly, the magnitude of DAN activation was generally similar across the five vapour exposures, apart from a small reduction between the first and second exposures (Supplementary Figure 3b). This activation pattern resembles that reported for MBON responses over repeated exposures to attractive odours ^27^, and indicates that DAN responses to vapour do not habituate, consistent with continued behavioural approach over successive intermittent trials (Supplementary Figure 1j).

To test whether DAN activity reflected upwind approach during the 1 s of water vapour exposure, we calculated a repeated measures linear fit between the vapour-evoked fluorescence and upwind displacement for each fly (using each 1 s vapour exposure from two protocol repeats per fly). Upwind displacement was positively correlated with γ4 and β′2 DAN activity and negatively correlated with γ5 activity (Supplementary Figure 3c), consistent with a prior report of γ4 DAN responses in flies tracking vinegar odour ^18^. To test whether this correlation depended on the direction vapour hits the fly, we analyzed data from water vapour exposures where the fly was stationary at vapour onset, to remove fluorescence associated with locomotion. These data revealed the magnitude of γ4 and β′2 DAN responses to be higher when the vapour hit the fly head on, versus from the side (Supplementary Figure 3d). Together, these data show that water vapour-evoked γ4 and β′2 DAN activity reflects upwind orientation relative to vapour and approach towards vapour.

In contrast, the magnitude of DAN activation by water vapour was not correlated with upwind displacement following vapour offset (data not shown), indicating that DAN activity in the presence of vapour does not predict future approach when vapour disappears. We therefore hypothesized that persistent approach is instead encoded by DAN activity within the 14 s off period. Aligning behaviour with imaging showed that locomotion is associated with elevated β′2, γ4 and γ5 DAN activity (Figure 3d).

We next performed a more detailed analysis of locomotion related DAN activity to decipher what information it contained. We first detected speed bouts using the fly’s forward/sideways locomotion in the 2-14 second period following water vapour offset (average of 8 bouts per protocol repeat, SD=3) and analyzed the corresponding bilateral changes in DAN activity during these bouts (an average 8 bouts (SD=3) detected per protocol repeat). Speed bouts were associated with a robust increase of γ4 and β′2 DAN activity, and with a weak response of γ5 DANs (Figure 3e). Cross correlation revealed speed bouts to be temporally coincident with both γ4 (Zolin et al., 2021) and β′2 DAN activity (Supplementary Figure 4a-b). Importantly, the behaviour to fluorescence signal cross correlation also shows that following our temporal data alignment (see methods) there is no measurable delay resulting from image acquisition of GCaMP8f fluorescence.

Our earlier experiments showed that upwind approach deviates during vapour offset when DANs are blocked (Figure 2). We reasoned that this likely results from a requirement for an increase in speed bout-evoked DAN activity when vapour is lost (i.e. a fluorescence increase dependent on stimulus history). We therefore tested for a contribution of stimulus history by comparing speed bout-evoked DAN activity following vapour offset with speed bout-evoked DAN activity in clean air (before the fly experienced water vapour). However, bout onset-triggered (Supplementary Figure 4c) and bout offset-triggered DAN activity (Supplementary Figure 4d) were similar irrespective of stimulus history. Running speed was also independent of stimulus history (Supplementary Figure 4 c-d). Although we observed a delayed decrease in speed bout onset-triggered β′2 DAN activity following water vapour exposure (Supplementary Figure 4c; 0.4-1 s window from speed bout onset; statistics not shown), less activity seems unlikely to account for loss of tracking when DANs are blocked. Therefore, speed bout-evoked DAN activity at this level does not appear to reflect stimulus history. Since knowledge of prior vapour is necessary to maintain tracking when it is lost, we reasoned that stimulus history must be otherwise represented in DAN activity.

We next tested whether DAN activity during speed bouts could represent directed locomotion by quantifying fly movement relative to wind direction. A repeated measures cross correlation was calculated between the magnitude of bilateral DAN activity and the fly’s rotation relative to upwind within each speed bout, using data extracted from speed bouts within the post-stimulus period and irrespective of stimulus history (Supplementary Figure 4e). This analysis showed γ4 and β′2, but not γ5, DAN activity to be higher during running bouts where flies turn upwind as compared to those when turning crosswind, suggesting that DAN activity includes a rotational component. We wondered whether these elevated DAN-type specific responses to upwind locomotion might resemble striatal dopaminergic neuron responses in mice, which reflect whether an optimal trajectory is being followed towards a learned goal location ^28^.

### Delayed γ4 and β′2 DAN activity encodes turns relative to allocentric wind direction

Prior evidence showed MBONs can influence steering ^14^, consistent with a prior suggestion that DAN type-specific modulation could alter steering through MBONs ^29^. Our alignment of imaging and behaviour data revealed flies to turn multiple times within an individual speed bout (Figure 3d; on average 10 ± 4 rotational bouts over protocol presentations, n=16), allowing us to test whether DAN activity relates to these steering manoeuvers.

To do this we detected upwind and crosswind turn bouts from the 2-14 s period following stimulus offset, irrespective of stimulus history (Figure 4a-b) and analyzed the associated turn-triggered bilateral DAN activity during the bouts (Figure 4c). Upwind turn-triggered activity was significantly greater than zero across all recorded DAN types, as found with speed bout evoked activity. In contrast, crosswind turns did not evoke significant activity in any DAN type, and upwind turning evoked stronger activity selectively in γ4 and β′2 DANs relative to crosswind turning. Flies made both upwind and crosswind turns of a similar angle (Figure 4b) and turns in either direction were associated with similar increases in speed (Supplementary Figure 5a-c). These data reveal that γ4 and β′2 DAN activity encodes turn direction relative to allocentric wind direction, with turns made upwind represented by a bilateral increase in DAN activity. To our knowledge, dopaminergic neurons have not previously been shown to encode orientation relative to a sensory cue during navigation.

**Figure 4.**
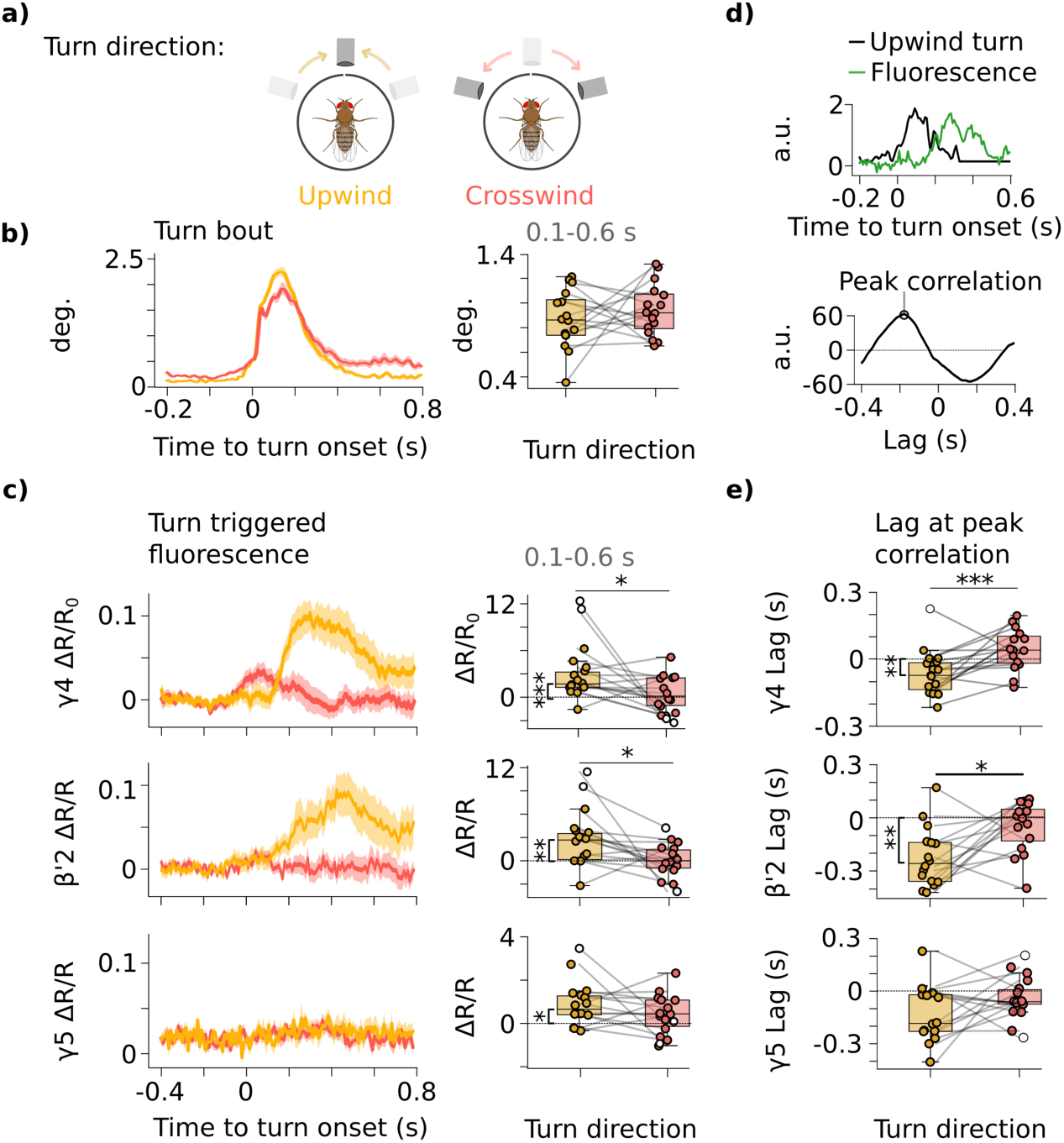
γ4 and β′2 DAN activity temporally follows goal-directed turning. **a,** Turns were separated according to allothetic turn direction (i.e. upwind or crosswind). Example movement of air port are shown, from previous (transparent) position to current (non-transparent) position along an upwind (yellow arrows) or crosswind (red arrows) path. **b,** Both upwind and crosswind turns have similar magnitude. **c,** Upwind turns, but not crosswind turns, evoke γ4, β′2 and γ5 DAN activity as measured relative to zero. γ4 and β′2 activity was significantly higher following upwind relative to crosswind turns. Traces plotted over time as mean ± SEM. Area under curve was measured using the 0.1-0.6 s duration following turn bout onset. **d,** top: Example upwind turn bout and corresponding γ4 DAN fluorescence (z-score normalized), and bottom: their cross correlogram showing the lag at peak correlation. **e,** γ4 and β′2 DAN activity lags behind upwind turning. Lags for upwind turns were significantly lower than zero and significantly lower than lags for crosswind turns (n=16 flies). Exact statistical values and comparisons are presented in Supplementary Information.

We next assessed whether DAN activity preceded or followed turns. Although our previous analysis revealed DAN activity to be temporally aligned to forward/sideways locomotion (Supplementary Figure 4a-b), aligning upwind turns with imaging revealed a clear temporal lag in DAN activity associated with turn onset (Figure 4d). A cross-correlation revealed γ4 DANs to be active 80 ms (median, IQR=0.13) after upwind turns and β′2 DAN activity by 150 ms (median, IQR=0.18) after upwind turns (Figure 4d-e). The same analysis of crosswind turns, that were not associated with significant DAN activity (Figure 4b), did not detect significant lagged activity, which controls against spurious correlation to signal noise. Delayed activity of dopaminergic neuron activity relative to action onset is indicative of performance evaluation ^30^. Taken together, these data reveal a role for DANs in evaluating turning manoeuvers relative to an allothetic wind cue, which supports the requirement of γ4 and β′2 DAN activity in maintaining upwind approach when vapour disappears.

### Lateralized γ4 DAN activity reflects egocentric turn direction

While memory-relevant computations involving mushroom body DANs are generally considered to be bilateral, it would seem logical that steering-related activity could be lateralized, like that of dopaminergic neurons directly associated with the compass network ^15^ or descending circuitry of locomotor control ^31^. To look for lateralized activity of γ4, γ5 and β′2 DANs during navigation, we extracted egocentric (i.e. clockwise and counterclockwise) turn-triggered DAN activity from both sides of the brain, irrespective of allocentric (i.e. upwind or crosswind) turn direction. Surprisingly, segmenting the data in this way revealed lateralized γ4 DAN activity, where turning evoked stronger activity in the mushroom body compartment ipsilateral to the egocentric turn direction (Figure 5a,b). In comparison, γ5 and β′2 DANs did not show lateralized activity (Supplementary Figure 6a,b).

**Figure 5.**
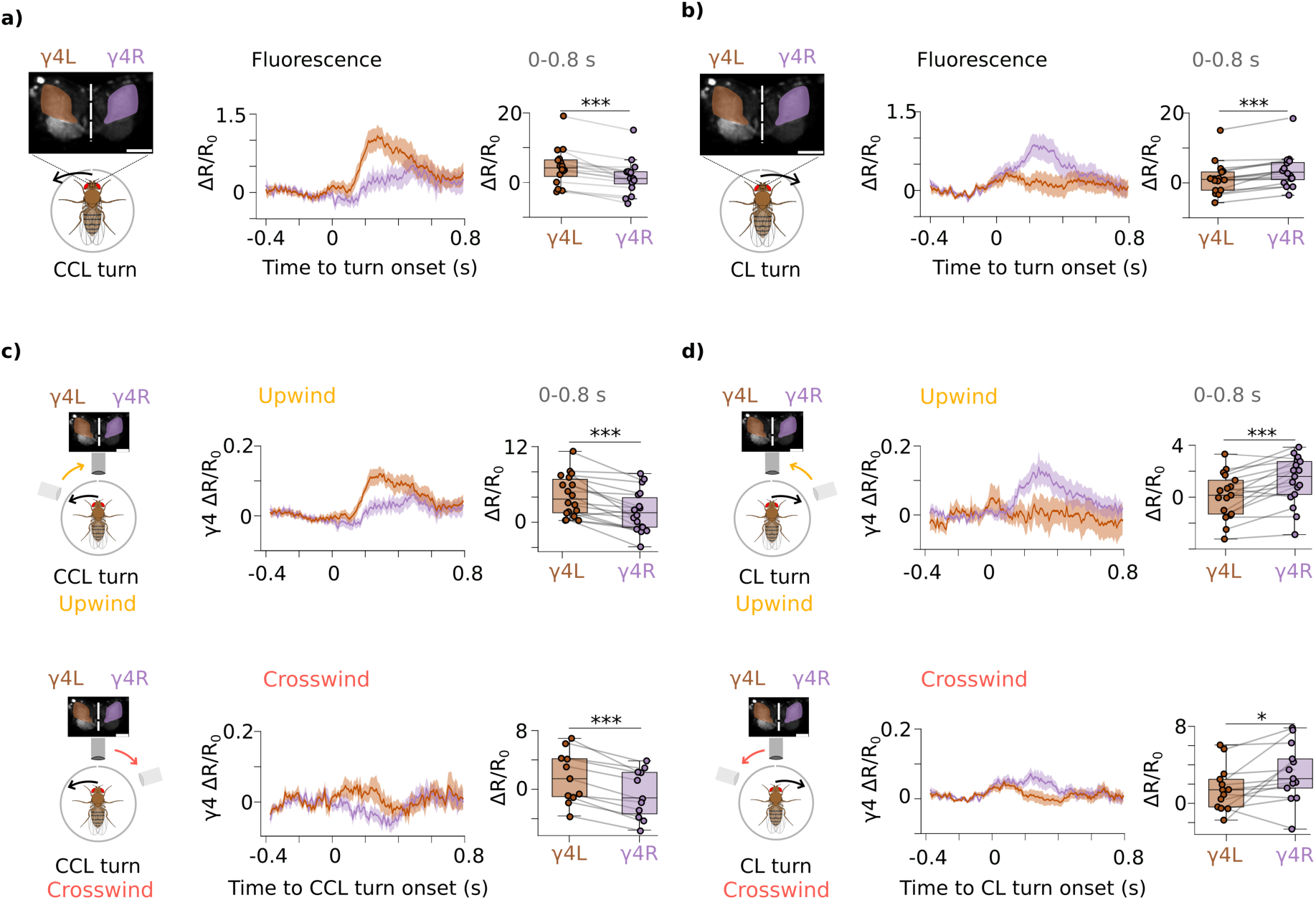
Ipsilateral γ4 DAN activity encodes egocentric turn direction. **a-b,** Turns were separated as counterclockwise, CCL (a), and clockwise, CL (b), and turn-triggered γ4 DAN activity was extracted from the left (γ4L; orange) and right (γ4R; purple) sides of the brain (cartoons indicate direction of turn that was used to group data). Boxplots show area under curve calculated from the 0-0.8 s duration from turn onset (each datapoint is the average area under curve for each fly; n=16 flies). **c-d,** γ4 DAN activity represents ipsilateral turning regardless of change in allothetic wind direction. Counterclockwise, (c) and clockwise (d) turns were further separated based on whether the turn was made towards upwind (yellow; top row) or crosswind (pink; bottom row). n=11-17 flies per pairwise group. Statistical comparisons made using two-sided paired Wilcoxon signed-rank test. All traces over time plotted as mean ± SEM. For brain images, solid line is scalebar (10µm) and dashed line the midline. Exact statistical values and comparisons are presented in Supplementary Information.

We hypothesized that a possible function of lateralized dopaminergic activity during navigation could be to keep track of egocentric turn direction, in which case ipsilateral turn-triggered γ4 activity should not be influenced by change in wind direction. To test this, turn-triggered DAN activity was compared between both sides of the brain for all four combinations of clockwise or counterclockwise turns made upwind or crosswind (Figure 5c,d). Consistent with our hypothesis, turns evoked greater ipsilateral γ4 DAN activity relative to the egocentric turn direction, regardless of change in wind direction. That is, turns to the left evoked DAN activity in the fly’s left hemisphere, and the right hemisphere for right turns.

We believe these data provide the first evidence of lateralized activity within dopaminergic neurons of the fly’s mushroom bodies. Finding that lateralized DAN activity represents locomotor turns that are independent of sensory cues led us to postulate that they may represent goal-directed turning in the absence of vapour.

### γ4 DAN activity encodes goal-directed turning

Our behaviour data thus far showed that brief exposures to water vapour evoked a persistent upwind approach in vapour absence (Figure 1), that was impaired when DANs were blocked (Figure 2). As already mentioned, DAN-dependent tracking in the absence of vapour suggests that history of vapour experience should also be represented in DAN activity. We therefore tested whether upwind or crosswind turn-triggered DAN activity reflected recent vapour exposure (i.e. stimulus history). To increase our analytical power, we accounted for laterality in turn-triggered γ4 DAN activity (Figure 5) by separately taking data from clockwise upwind turns from the right hemisphere and counterclockwise upwind turns from the left hemisphere (Figure 6a-b) and combined these responses as ‘upwind’. This analysis revealed vapour history to be represented as elevation of γ4 DAN activity when flies turned upwind, during periods when vapour was off. A similar analysis revealed no effect of stimulus history on crosswind turn-triggered γ4 DAN activity (Supplementary Figure 7a-b).

**Figure 6.**
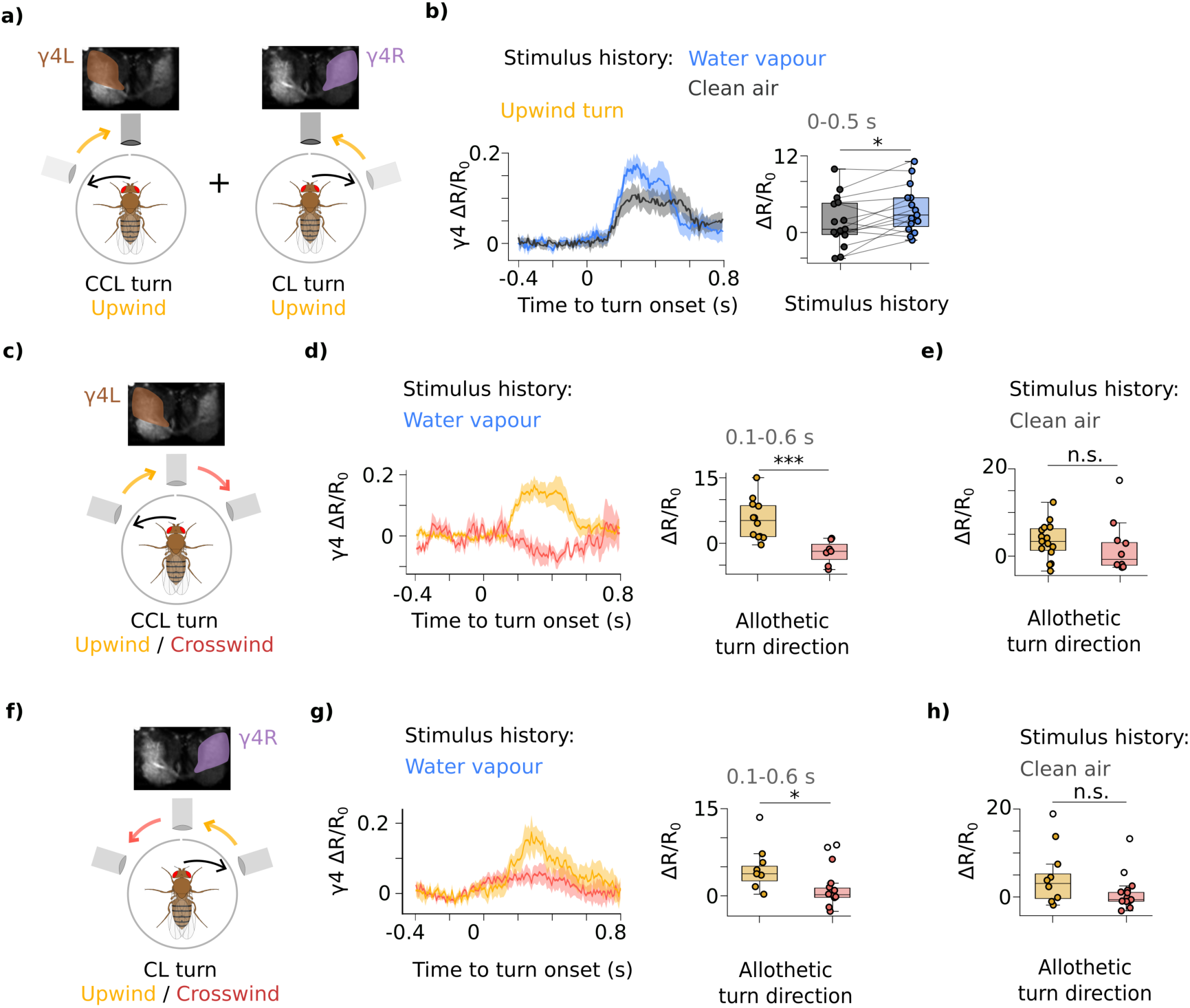
Ipsilateral γ4 DAN activity encodes goal-directed turning. **a,** To improve signal strength, ipsilateral turn-triggered γ4 DAN activity was obtained by combining activity from left hemisphere during counterclockwise (CCL) turns with activity from right hemisphere during clockwise (CL) turns, only using turns that oriented the fly upwind (yellow arrows). **b**, Upwind turn-triggered γ4 DAN activity was greater following water vapour exposure relative to turns in clean air. Boxplots show area under curve measured in 0-0.5 s from turn onset (n=15-16 flies per pairwise group; two-sided paired Wilcoxon signed-rank test). **c-h,** γ4 DANs encode stimulus history as an ipsilateral increase in upwind turn-triggered activity. Ipsilateral γ4 DAN activity was grouped by egocentric turn direction, with left side brain activity taken for counterclockwise turns (c-e) and right side brain activity taken for clockwise turns (f-h). γ4 DAN activity was greater ipsilaterally when the turn was made upwind (yellow), relative to crosswind (pink), following water vapour exposure (e,h) but not in clean air (f,i). n=7-12 flies per group. Statistical comparisons made using a Wilcoxon rank-sum test (Mann–Whitney U test). Cartoons in c,f show egocentric turn direction (black arrow) that orient the fly towards either upwind (yellow arrow) or crosswind (pink arrow). All traces plotted over time represented as mean ± SEM. Exact statistical values and comparisons are presented in Supplementary Information.

To test whether stimulus history is specifically represented in γ4 DAN activity, we also analyzed turn-triggered bilateral responses of β′2 and γ5 DANs. In contrast to γ4 DANs, γ5 and β′2 DANs showed no evidence of representing stimulus history (Supplementary Figure 7a). The γ4 DANs still showed representation of stimulus history when bilateral responses were quantified, although the history contribution was less apparent as compared to analyzing ipsilateral activity. The increase in upwind turn-triggered γ4 DAN activity following vapour exposure does not result from increased forward locomotion, as flies travelled a similar amount following upwind turn onset regardless of vapour history (Supplementary Figure 7d). These data therefore reveal that γ4 DAN activity uniquely reflects goal-directed turning.

Our previous analysis revealed ipsilateral activity in γ4 DANs encodes whether the fly turned left or right (Figure 5). Finding that γ4 DAN activity reflects appropriate turn direction led us to predict that lateralized activity in these DANs could further encode the direction in which the appropriate turn was made. If this is the case, then ipsilateral γ4 DAN activity would not only keep track of left and right turns, but activity in each hemisphere should further encode whether the turn orients the fly to face upwind where water vapour was previously encountered. In other words, activity in each hemisphere ipsilateral to the egocentric (clockwise or counterclockwise) turn direction should encode allocentric (upwind versus crosswind) turn direction depending on stimulus history (water vapour or clean air). To potentially visualize this, we extracted data following vapour offset, and grouped turn-triggered γ4 DAN activity collected from flies making an upwind or crosswind turn according to the hemisphere ipsilateral to turn direction selected (i.e. the left hemisphere for counterclockwise turns and the right hemisphere for clockwise turns; Figure 6c-h). Confirming our prediction, appropriate counterclockwise (Figure 6c,d) or clockwise (Figure 6f,g) turns towards upwind, that bring the fly in the direction where vapour was encountered, evoked higher ipsilateral γ4 DAN activity as compared to inappropriate crosswind turns. In comparison, turns in clean air (prior to vapour exposure) had no ipsilateral representation in the γ4 DANs that reflect upwind or crosswind turns (Figure 6e,h).

Taken together, our data reveal novel roles for subsets of mushroom body DANs in representing different elements of goal tracking. β′2 and γ4 DANs evaluate turn manoeuvers relative to an allocentric wind direction, with turns that bring the fly to face upwind evoking delayed β′2 and γ4 DAN activity. Ispilateral activity in the γ4 DANs further represents egocentric turn direction, allowing the fly to keep track of which direction it turned regardless of wind fluctuations. And finally, we found evidence for vapour history to be represented by ipsilateral γ4 DAN activity, that reflects which direction an appropriate turn was made while searching for water. It will be interesting to determine whether distinct signals attributed to γ4 DANs involve the same individual neurons, or parallel subtypes ^29^.

## Discussion

Keeping track of actions and evaluating outcomes is essential for biological and artificial neural networks to determine progress towards a goal. During reinforcement learning, dopaminergic neurons perform these goal-directed computations by representing action relative to a learned reward location ^28,32,33^. Similar computations could be employed during olfactory navigation in search of resources ^34–36^, which requires representing actions relative to environmental cues that predict reward direction. We provide the first evidence that lateralized activity in mushroom body γ4 dopaminergic neurons follows turns made in the appropriate direction, allowing flies to stay on track towards a goal when cues are intermittent.

Despite exclusively studying approach to water vapour, we predict that the MB will similarly maintain search towards other fluctuating volatiles that have innate or learned positive valence. Supporting this expectation, γ4 and β′2 DANs preferentially respond to innately appetitive odours ^27^ and their activity can reinforce learning by inducing plasticity of KC-MBON connections. We propose such plasticity, perhaps between hygrosensory α′β′ KCs and connected MBONs, will be how transient memory of water vapour is represented. Since MBONs, including those modulated by γ4 and β′2 DANs, directly synapse onto central complex neurons ^29^, lateralized evaluative activity could be transferred from the MB to central complex and downstream descending neurons controlling turning ^11,15,31,37^. Although individual γ4 and β′2 DANs actually project bilaterally, an ipsilateral steering signal can arise from their denser ipsilateral projections, or asymmetric modulation of arbour activity.

Importantly, our data show that goal-directed turning evokes delayed activity in the γ4 DANs. This DAN turn-evoked activity could arise from recurrence from post-turn signals in the central complex ^13^ and pathways that compute changes in wind direction ^38^, although direct central complex afference to DANs is limited and does not include connectivity to γ4 and β′2 DANs ^29,39^. Temporal delays are a feature of phasic dopamine signals that encode prediction errors during reinforcement learning. Finding that fly dopaminergic neurons provide moment to moment comment on the appropriateness of selected steering directions is reminiscent of those in the songbird, which evaluate vocal performance ^30^. The mushroom body therefore provides the capacity to evaluate how a completed turn compares against expectation, complementing the navigational computations performed within the central complex. Further study in the fly has potential to reveal conserved circuit computations underlying performance evaluation.

## Methods

### Fly Strains

The wild-type *D.melanogaster* strain was Canton-Special ^40^. UAS-*Shi*^ts1^(X;3) flies are described ^41^. R48B04-GAL4 flies were previously characterized ^42^. For two-photon imaging experiments *20XUAS-IVS::jGCaMP8f* ^43^ and *UAS-myr::tdTomato* ^44^ were expressed under the control of R48B04-GAL4.

### Fly Husbandry

All strains were maintained at 22 °C and 60 % humidity in a 12:12 h light:dark cycle with light provided between 8 AM and 8 PM. For all behavioural experiments, flies were reared on yellow cornmeal agar food containing deionized water, 7.2 g l−1 agar (Fisher Scientific), 25 g l−1 autolysed yeast extract (Brian Drewitt), 47.3 g l−1 cornmeal (Brian Drewitt), 100 g l−1 dextrose (d-glucose anhydrous, Fisher Scientific), 2.2 g l−1 tegosept (methyl 4-hydroxybenzoate) (Sigma-Aldrich), and 8.4 ml l−1 ethanol (Sigma-Aldrich). All cornmeal agar food was prepared by boiling, not autoclaving.

### Water and Food deprivation

For dehydration, approximately 30 flies per vial were water deprived by housing them for 18-24 h at 21-25 °C at 30 % relative humidity in a climate chamber (Memmert HPP110 eco). The exact duration and temperature was varied dependent on genotype, as some genotypes dehydrated quicker than others. For starvation, flies were food deprived for 20-26 h in a 25 ml vial containing a 2 x 3 cm piece of filter paper with 1 % agar at the base. Vials were stored at 22 °C throughout starvation. Satiated flies were provided with *ad libitum* access to food and water.

### T-maze behaviour

T-maze behaviour in Supplementary Figure 1c was performed in either satiated, hungry or thirsty wild-type flies. Groups of 100 flies were given 2 min to choose in darkness between two tubes, one that carried a dry air flow and a ‘vapour’ tube that carried air bubbled through distilled water.

### Measuring locomotor behaviour

The air-supported trackball setup was adapted from Seelig et al. (2010) ^45^. A camera (Point-Grey, GS3-U3-23S6M-C NIR) was used to acquire images of dots drawn on a 6.35 mm diameter, 0.6 gram aluminium ball (McMaster Carr, 1089T21) at 70 frames per second. These images were used by the program Fictrac ^46^ to reconstruct the walking trajectory of the fly. Positive/negative values associated with rotatation around the fly x,y, and z axes represent right/left, forward/backward and clockwise/counterclockwise steps, respectively. For each frame acquired by Fictrac, the delta (change since previous frame) x,y,z coordinates were streamed into a custom written script in python (v 3.10). Each frame was associated with a timestamp that was used to align fly behaviour with data from other devices (e.g. stimulus on/offsets, imaging data, arduino).

### Tethering flies

For pin-tethering, flies were first cold anesthetized by placing them on ice for 30 seconds. They were then transferred to a cold plate where UV light (Thorlabs UV Curing System, CS2010) was used to cure a droplet of UV glue (Norland optical Adhesive 68) between the tip of a 0.06 mm diameter insect pin and the thorax of the fly. Flies were then allowed to recover for 20 minutes by giving them a small ball of dried sugar paper.

For head-tethering, flies were first pin-tethered as described above. While still cold anaesthetized, a droplet of UV glue was placed between the head and thorax, and the head was angled downwards by approximately 45 degrees before curing the glue. The pin-tethered fly was then positioned below a 3D printed platform with a 0.45 mm hole cut out for the head. UV glue was then used to glue the eyes and thorax to the platform. A bridge made of bee wax was then built on the top of the thorax using a wax melter (Almore).

Flies were then positioned on the ball using a 3-axis manipulator (Märzhäuser MM33), with the help of a top-down camera (Thorlabs CMOS CS165MU) and a sideview camera (PointGrey GS3-U3-23S6MC).

### Shibire block on the ball

After head-tethering, a 50 μL drop of saline (103 mM NaCl, 3 mM KCl, 4 mM MgCl_2_, 1.5 mM CaCl_2_,5 mM HEPES-NaOH,10 mM trehalose, 10 mM glucose, 26 mM NaHCO₃, 1mM NaH₂PO₄, pH 7.2, 275 mOsm kg−1) was placed on top of the fly’s head, without opening the head capsule. A wire attached to a temperature controller (Thorlabs TC300B) was immersed into a drop of saline on top of the fly’s head. A thermocouple attached to the wire read out the actual temperature of the wire. For restrictive temperature, the wire was set to 34 °C for 20 min before stimulus exposures. Saline was kept at 34 °C to replenish the drop on the fly’s head as required. For permissive temperature controls, the wire was set to 22 °C.

### Stimulus delivery

The mcculw (v1.0.0) python package was used to adjust flow controllers (Sensirion SFC5500) via a data acquisition device (Measurement Computing USB-1208LS) and analog output device (Measurement Computing USB 3103) that maintained air flow at a rate of 0.2 litres per minute. The python package nidaqmx (v 0.8.0) was used to control an input/output module (National Instruments USB-6509) that controlled a custom-built spike and hold circuit (Shang et al., 2007) to toggle valves (Lee Company) that directed air flow.

A constant 0.2 liter per minute (LPM) clean airflow was directed through odourless tubing (Cole-Parmer, Versilon) that terminated in a 1.4 mm diameter cylindrical stimulus port (McMaster Carr, <u>5560K74)</u> positioned ∼4 mm from the head of the fly. Either clean air or water vapour stimulations were delivered in open loop, by replacing the constant airflow by air bubbled through mineral oil or purified water (Milli-Q^®^) respectively, such that the flow rate always remained at 0.2 LPM. The air carrying the stimulus was introduced via a fast switching three-way valve (Lee Company) placed ∼30 cm away from the fly. Passing the air through a flow sensor (Honeywell, AWM3300V) shows minimal pressure fluctuations were associated with opening and closing of the valves. Stimulus onset time was shifted by ∼0.5 s to account for the time taken for the stimulus to travel the length of tubing from the three-way valve to the fly, which could be measured by a photoionization detector (Aurora miniPID). The miniPID was normally positioned behind the fly and suctioned air at 0.8 LPM.

For figure 3c, observed fluorescence during clean air exposure was due to the minor pressure fluctuation associated with toggling the air valves. This pressure fluctuation explains the bi-phasic appearance in the 1 s stimulus exposure in both the clean air and water vapour exposures; the onset and offset timepoints of the stimulus are right-shifted by 0.5 seconds to accommodate for the time taken for the stimulus to travel the length of tubing from the valve to the fly, which coincides with the dip in fluorescence.

The stimulus port was connected to a servo (Clearpath CPM-SDSK-3411S-ELN) set at 800 step resolution, 10,000 rotational velocity and 200,000 rotational acceleration. Step signals were sent by an Arduino Mega 2560 running at ∼120 loops per second, using the ClearPathStepGen.cpp library. Fly heading was monitored in real time by Fictrac and was yoked to the stimulus port in closed loop, such that the fly could control its angle of travel within the air stream. Rotations were calculated in 1 ° steps, such that the fly had to turn at least 1 ° to trigger a corresponding rotation of the stimulus port, with –75° to 75° limits from head-on.

### Behaviour analysis

The servo position was acquired in real time at 120 Hz (loop rate of the Arduino), and was first resampled to match the behaviour (captured at 70Hz; see above). These data were then interpolated to account for dropped frames during acquisition. The fly’s x/y/z coordinates (measured by Fictrac) were then transformed to obtain the walking trajectory relative to the wind direction (measured by the Arduino).

Fly speed was calculated from the change in xy coordinates (i.e. forward/sideways position) between successive timepoints measured by Fictrac at 70 Hz. In Figure 1e,h, rotation was calculated as the change in fly z rotation between successive timepoints and converted to degrees per second. A protocol repeat was only used for analysis if the fly walked an average 3 mm/s over the protocol duration.

To measure angle of travel over a given time, a linear fit was made using the x-axis as upwind or crosswind depending on which had the most spread in data, and weights as the distance between each datapoint (implemented via python numpy.polyfit). The angle relative to upwind was then used as the angle of travel for that duration of time. Angle of travel was only measured if the fly travelled at a speed of 3 mm/s over the duration at which the angle was being calculated.

### Two-photon in vivo calcium imaging

Flies up to 10 d old were water-deprived for 18-24 h. After head-tethering a fly, the head capsule was opened under hyper-osmotic saline (120 mM NaCl, 3 mM KCl, 4 mM MgCl_2_, 1.5 mM CaCl_2_,5 mM HEPES-NaOH, 10mM trehalose, 10mM glucose, 26 mM NaHCO₃, 1mM NaH₂PO₄, pH 7.4, 320 mOsm kg−1). Two-photon imaging was performed using a multiphoton imaging system (Sutter MOM) with a 20x, 1 NA water-immersion objective (Olympus). A green (525/30, Semrock) and red (610/75, Chroma) filter were followed by GaAsP PMTs (Hammamatsu) to detect GCaMP8f and tdTomato fluorescence, respectively. Fluorescence was excited at 910 nm, repetition rate 80 Hz, pulse duration ∼70 fs, using a Ti:sapphire laser (Mai Tai DeepSee). Images (128 × 128 pixels) were acquired at ∼110 Hz, controlled by ScanImage (2022.1.0) software via MATLAB (MathWorks, release 2022a). The Scanimage DataRecorder was used to acquire TTL pulses from the Fictrac camera, stimulus exposures and image frame acquisitions. These TTL pulses were used to temporally align all recorded devices.

### Calcium imaging analysis

Acquired .tif files from imaging were first motion corrected using non-rigid motion correction ^47^ implemented in python. Fiji (v 1.54t) ^48^ was used to demarcate the anatomically distinct β′2, γ4 and γ5 ROIs manually for R48B04-driven expression of GCaMP6f and tdTomato. For each ROI, the mean intensity of each frame was extracted for each of the two channels (green i.e. GCaMP8f and red i.e tdTomato) recorded. A control region of a similar area was subtracted for background correction. To control for z-axis motion related artefacts, the mean intensity of each GCaMP6f frame for an ROI was divided by the mean intensity of its corresponding tdTomato frame. To correct for photobleaching, the signal was then divided by a mono-exponential fit made to the signal. This yielded the ratiometric fluorescence trace R, defined as the photobleach-corrected ratio of the GCaMP8f to tdTomato signal for an ROI (e.g. Figure 3d).

R was normalized by calculating ΔR/R_0_, with ΔR = R – R_0_ and R_0_ calculated as the average fluorescence prior to stimulus or behaviour onset. However, for Supplementary Figure 3b, R_0_ was calculated as the 40^th^ percentile for each 1 s on 14 s off duration since there was no separate baseline period for each vapour exposure.

### Behaviour-triggered fluorescence

Behaviour was first up-sampled to match the 110 Hz image acquisition rate. Bouts of speed or turns were found by using thresholds (Table 1) to detect their onset and offset timepoints. To detect turn bout, we used the servo rotation as it functioned as an additional lowpass filter that removed smaller heading changes associated with noise in Fictrac image processing that did not produce a change in wind direction (see above for how fly rotation was converted to servo rotation).

**Table 1.**

|  | speed | rotation |
| --- | --- | --- |
| lowpass | 5 | 10 |
| Upper threshold | 7 mm/s | 1 degree |
| Lower threshold | 2 mm/s | 0.4 degrees |
| width | 0.5 seconds | 0.2 seconds |
| distance | 0.5 seconds | 0.2 seconds |

To find the onset of a bout, data was first lowpass filtered using a second order butterworth filter to keep only lowpass frequencies, implemented via python scipy.signal. The timepoints where the measured behaviour passed an upper threshold were then detected. The offset timepoint was then found as the timepoint where the measured behaviour averaged over a distance rolling window fell below a lower threshold. A bout was only kept if it had a threshold width and if it was separated by a threshold distance from its neighbours. At least 2 bouts had to be detected during a stimulus exposure to consider a protocol repeat for analysis. The thresholds used for bout detection for different behavioural variables are as follows:

The onset of these behaviour bouts were then used to extract bout-triggered ratiometric fluorescence.

To obtain a cross correlation between the behavioural bouts and corresponding bout-triggered fluorescence, all waveforms were first z-score normalized. Correlation was implemented using the scipy.signal.correlate function in python, and the output correlation had the same number of datapoints as the input waveforms. The lag was obtained from each cross-correlation as the timepoint at peak correlation. The average lag was then obtained for each fly.

## Statistics

Each datapoint representing the average value for one fly (i.e. n=1) was obtained by first averaging within each protocol, and then averaging across the protocol repeats per fly. For comparisons between two paired groups, a two-sided paired Wilcoxon signed-rank test was used. For comparisons between two independent groups, a two-sided Wilcoxon rank-sum test (Mann–Whitney U test) was used. Differences among three or more independent groups were assessed using the Kruskal–Wallis test, and when significant, comparisons were performed using Dunn’s test with Bonferroni correction for multiple testing. Comparisons among three or more paired conditions were performed using the Friedman test, and when significant, post hoc pairwise comparisons were conducted using the Nemenyi test. Statistical significance is indicated in figures as P < 0.05 (*), P < 0.01 (**) and P< 0.001 (***).

A repeated measures correlation was implemented via the pingouin package (v 0.5.5) in python, using a linear fit for each fly with each fit having equivalent slope across flies. The coefficient r indicates the strength of correlation, and ranges from –1 to 1.

## Data availability

The behaviour and imaging data that support the findings of this study are available in Zenodo with the identifier 10.5281/zenodo.22896845.

## Acknowledgements

The authors thank L. Duquenoy and A. Cook for comments on the manuscript, and other Waddell lab members for discussion. A.M and SW were funded by a Wellcome Principal Research Fellowship (200846), a Wellcome Discovery Award (225192), an ERC Advanced Grant (789274), and Wellcome Collaborative Awards (203261 and 209235) to S. W.

## Author contributions

Designed research A. M., S. W., Built behavioural and microscope set up A.M. Performed research A. M., Analysed data A. M., Resources S. W., Writing A. M., S. W. Supervision S. W., Funding Acquisition S.W.

