## Supplementary Figures for "Lateralized dopaminergic neuron activity encodes goal-directed turning"

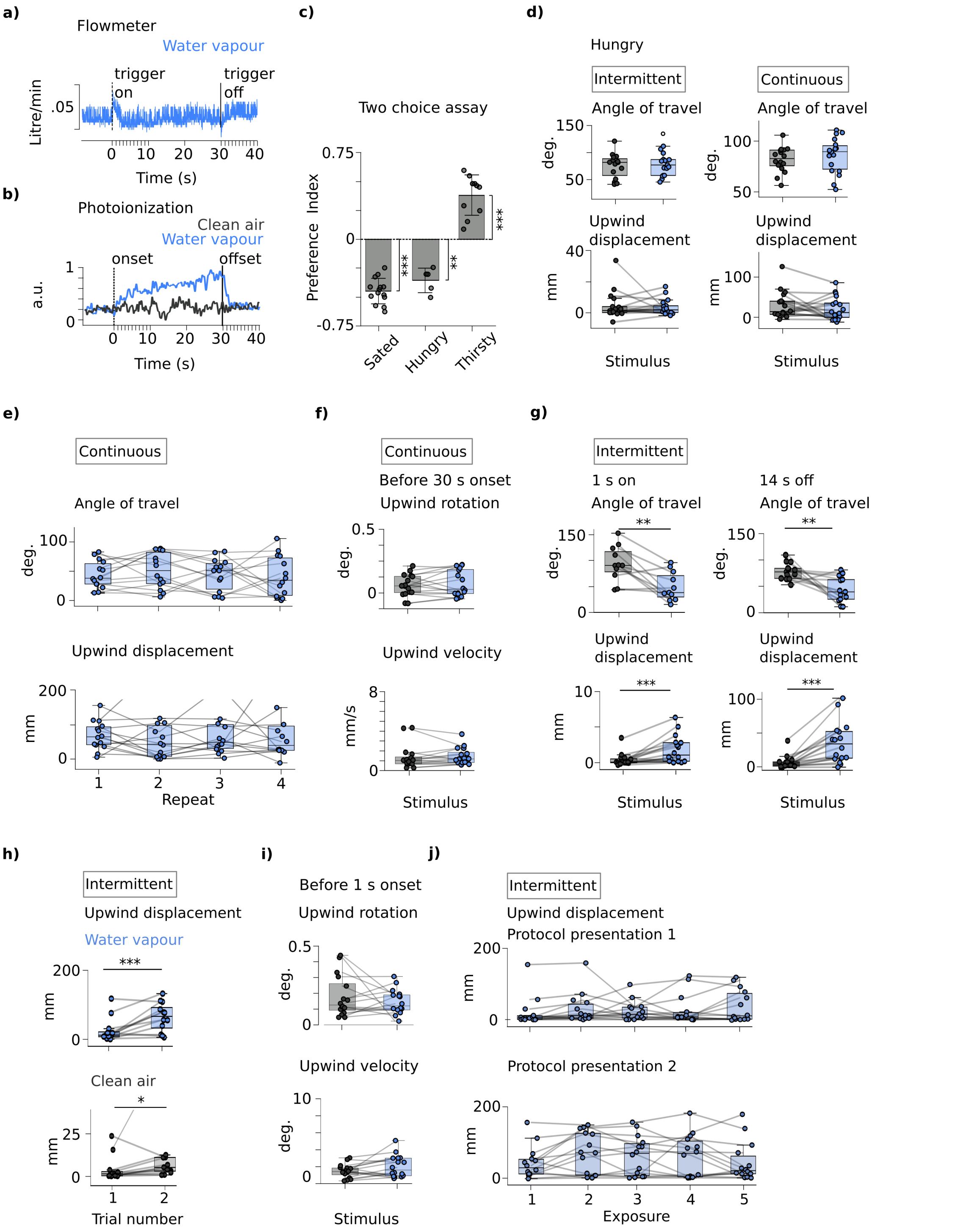


**Supplementary Figure 1. Vapour approach depends on internal state and stimulus protocol. a,** A flowmeter reveals minimal airflow turbulence when toggling air flow valves. **b,** A photoionization detector shows vapour reaches the fly at the onset timepoint and is gone within 2 s following offset. **c,** Sated flies (n=14) and hungry flies (n=5) avoid water vapour whereas thirsty flies (n=10) approach it in a two-choice assay. Positive Preference Index values indicate approach, with significance from 0 measured using a pairwise Mann-Whitney

U test. **d,** Hungry flies tethered to the trackball do not approach water vapour in either continuous or intermittent protocols. (n=15-17 flies per group; a two sided Mann-Whitney U test was used for angle of travel since few flies travelled at the threshold speed of 3 mm/s; see methods). **e,** There is no change in angle of travel (n=14 flies) or upwind displacement (n=13 flies) over four repeats of the continuous protocol (two-sided Friedman pairwise test; datapoints from two flies above y-axis limit not shown). **f,** Behaviour prior to vapour stimulus is similar to behaviour prior to clean air stimulus in the continuous protocol (measured over a 5 s duration; n=15-16 flies per pairwise group). g**,** In the intermittent protocol, approach is observed during the 1 s vapour on (left column) and in the 14 s following vapour offset (right column). n=13-16 flies per pairwise group. **f,** Approach towards vapour increases across the two repeats of the intermittent protocol (n=10-12 flies per pairwise group; datapoint from one fly above y-axis limit not shown). **g,** For the intermittent protocol, behaviour before first vapour exposure is similar to behaviour before first clean air exposure (measured over a 5 s duration; n=15-16 flies per pairwise group). **h,** For intermittent protocol, behaviour across exposures within each protocol repeat does not change (n=14 flies; two-sided Friedman pairwise test). Unless otherwise stated, comparisons between two groups made using a two-sided paired Wilcoxon signed-rank test. Exact statistical values and comparisons are presented in Supplementary Information.

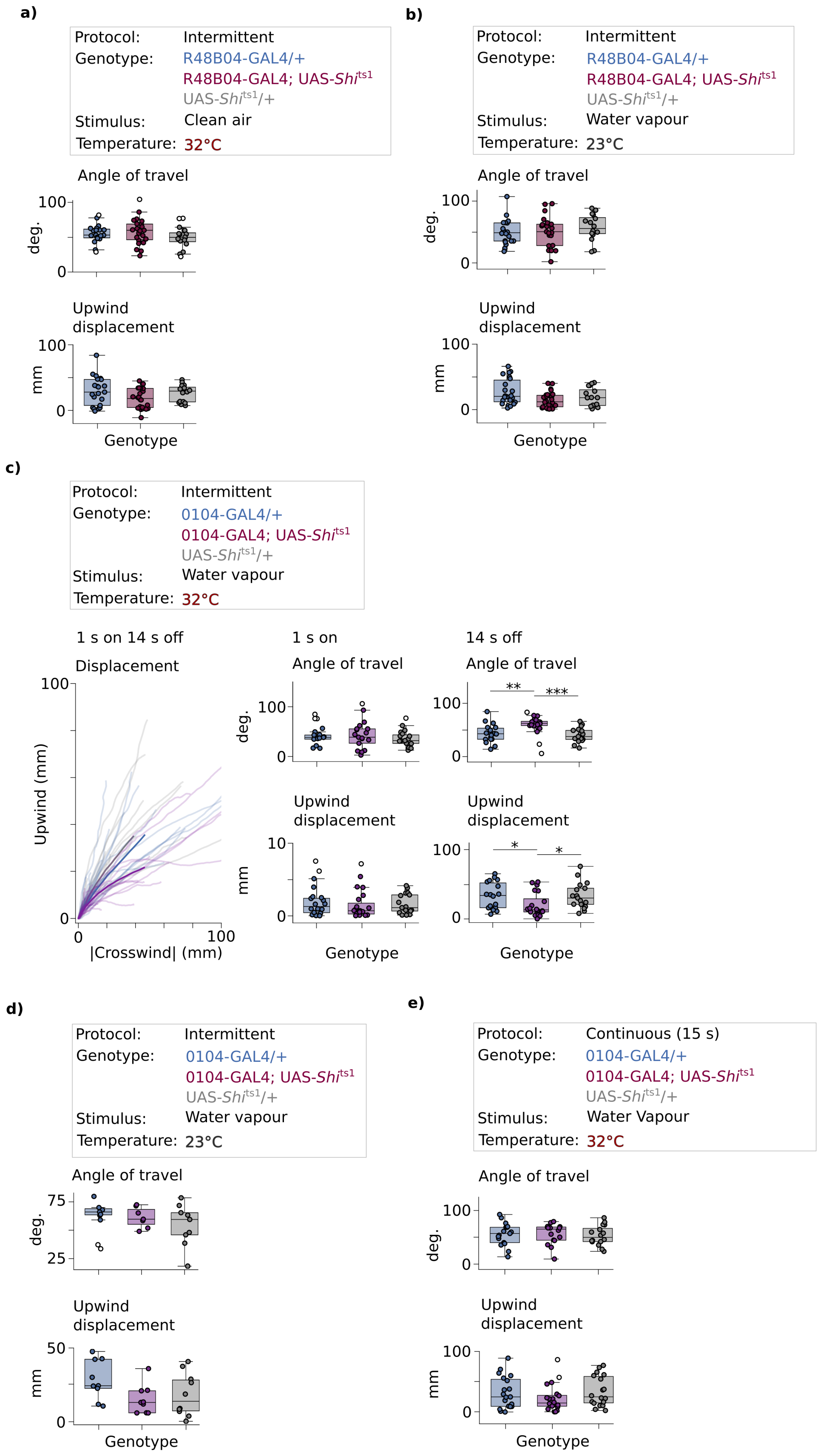


**Supplementary Figure 2. Similar impairment in post-vapour approach is observed with 0104-GAL4 neuronblock. a-b,** UAS-*Shi*$^{ts}$ mediated block of0104-GAL4 neurons does not impair a) behaviour in clean air at restrictive 32 ˚C (n=19-24 flies per group) or b) approach to water vapour at the permissive 23 ˚C (n=16-26 flies per group). **c-e,** UAS-*Shi*$^{ts1}$ block of 0104-GAL4 neurons, which partially overlaps with γ4 DANs and fully overlaps with β′2 DANs labeled by R48B04-GAL4, only impairs approach during vapour offset in the intermittent protocol (behaviour averaged across 5 exposures and two protocol repeats per fly; n=18-20 flies per group). **d-e,** There is no impairment in vapour approach d) at permissive 23 ˚C in the intermittent protocol (n=9-10 flies per group) or in e) at restrictive 32 ˚C in the continuous protocol (n=17-21 flies per group).  Hollow dots indicate outliers that are > 1.5 * inter-quartile range. Statistical comparisons made using a Kruskal-Wallis test followed by Dunn's multiple comparisons test with Bonferroni correction.

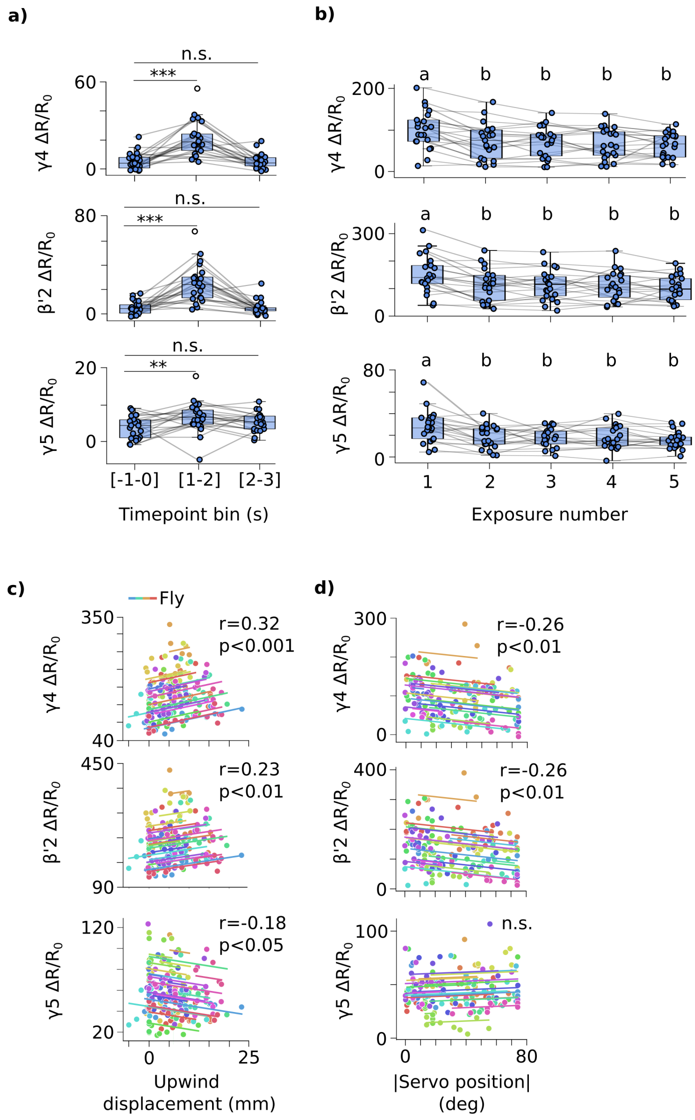


**Supplementary Figure 3. Vapour-evoked DAN activity correlates with upwind orientation and approach. a,** Water vapour-evoked activity in γ4, β′2 and γ5 DANs returns to baseline after two seconds following stimulus offset (n=22 flies per pairwise group). **b,** Vapour-evoked DAN activity resists habituation, with only first exposure evoking higher activity in all recorded DANs (n=22 flies). Different letters above bars indicate activity within exposures that are significantly different. Statistical tests in a-b performed with a two-sided Friedman pairwise test with a Neymeni post-hoc. **c,** Upwind displacement is positively correlated with vapour-evoked activity in all three DANs, as shown by a repeated measures correlation calculated between the area under curve for each 1 s water vapour exposure and the corresponding upwind displacement during that 1 s (n=22 flies). **d,** Direction that vapour hits the fly at stimulus onset is correlated to activity in γ4 and β′2 DANs, with higher activity when the fly faces directly upwind at 0 ° (using protocol repeats where the fly did not move, so to remove contribution of locomotion evoked fluorescence; n=22 flies). Exact statistical values and comparisons are presented in Supplementary Information.

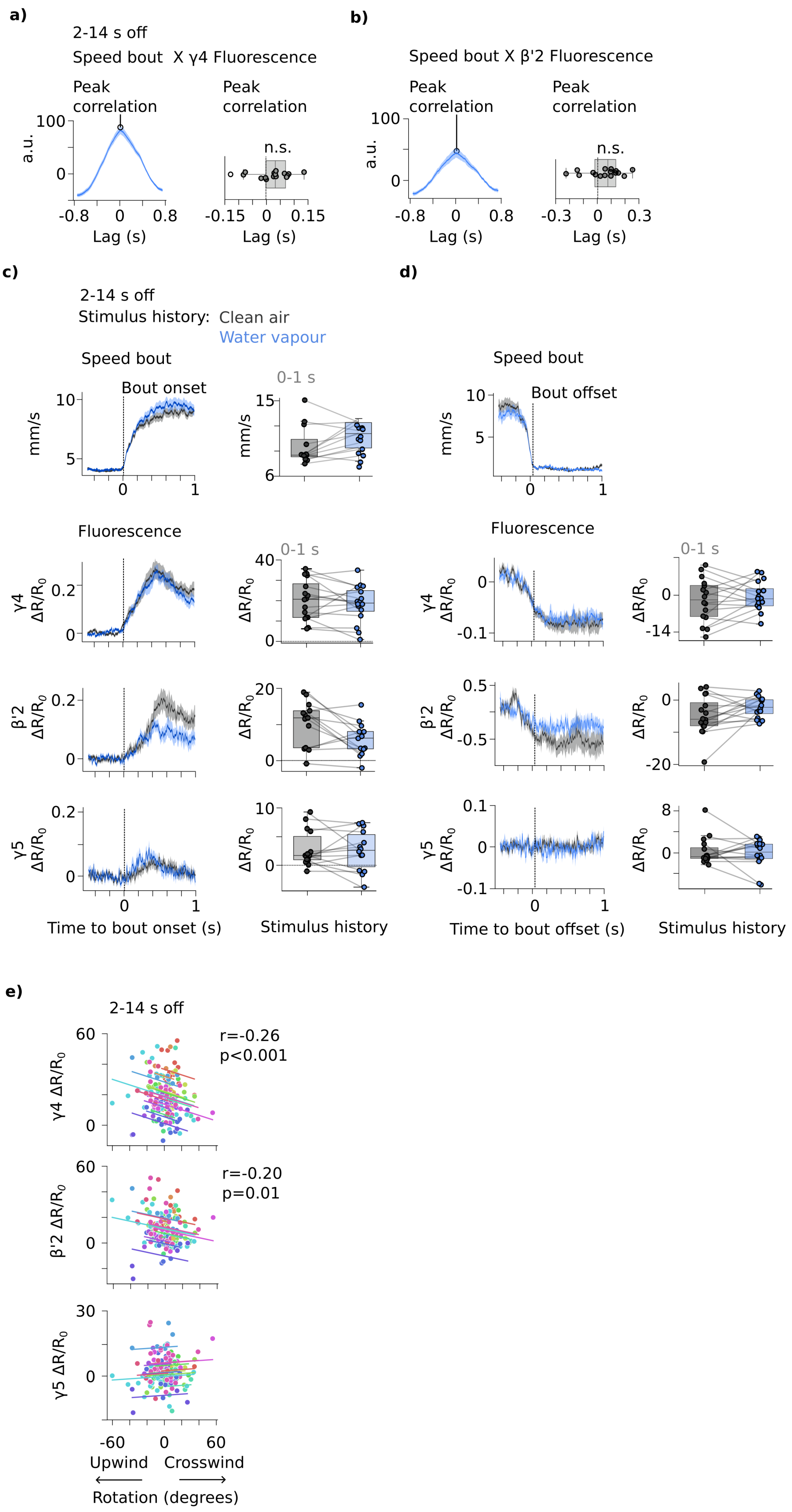


**Supplementary Figure 4. Speed bout-triggered DAN activity correlates with upwind locomotion. a-b,** Speed bouts temporally coincide with speed bout-triggered γ4 (a) and β′2 (b) DAN activity. Average cross-correlogram was obtained in the 2-14 s duration following water vapour offset (after averaging across all correlations from each fly). Peak correlation indicated for the averaged correlation waveforms (n=16 flies). Box-plot shows average temporal lag at peak correlation for each fly. Significance from zero measured using two-sided paired Wilcoxon signed-rank test. **c-d,** Speed bout-triggered fluorescence does not vary with stimulus history. Speed bout onset- (c) and offset- (d) triggered fluorescence in clean air (stimulus history=clean air) and following water vapour offset (stimulus history=water vapour), with area under curve measured in the 0-1 s duration (n=16 flies). **e)** Speed bout onset-triggered $\gamma 4$ and $\beta' 2$ activity is correlated with upwind orientation, such that running upwind evokes stronger fluorescence. Repeated measures correlation performed using fluorescence area under curve and sum total rotation measured from 0-1 s following speed bout onset (n=16 flies).

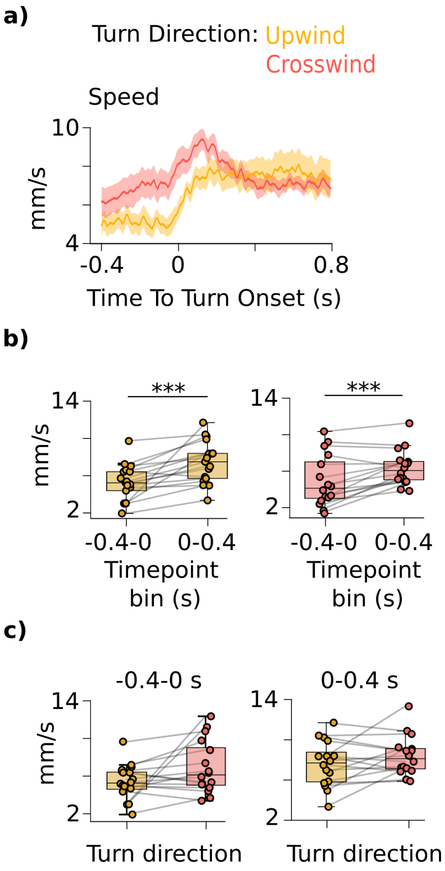


**Supplementary Figure 5. Walking speed does not drive the difference in upwind and crosswind turn-triggered fluorescence. a)** Upwind and crosswind turn-triggered speed. **b-c)** Both upwind and crosswind turns are associated with an increase in speed (b) and there is no difference in upwind or crosswind turn-triggered speed either before or after turn onset (c). Traces plotted over time as mean ± SEM. n=16 flies. Exact statistical values and comparisons are presented in Supplementary Information.

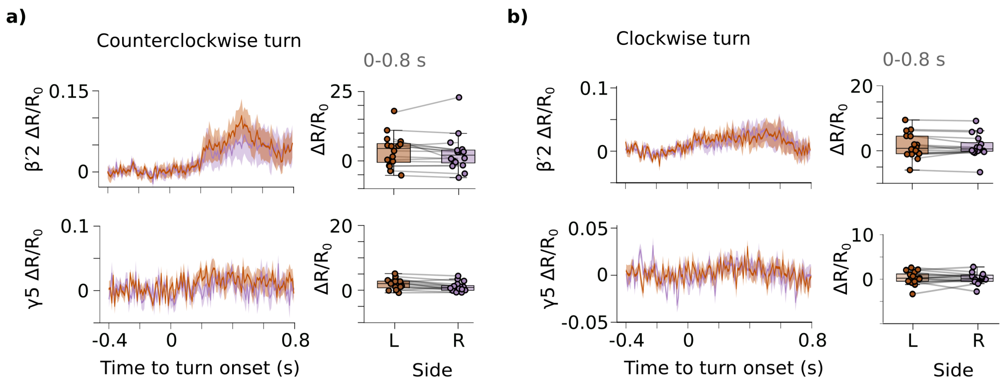


**Supplementary Figure 6. β′2 and γ5 DAN activity does not reflect egocentric turn direction. a-b,** Counterclockwise (a) and clockwise (b) turn-triggered β′2 and γ5 DAN activity reveals no difference between the left (orange) and right (purple) sides of the brain. Boxplots show area under curve measured from the 0–0.8 s window following turn onset. Statistical comparisons were made using two-sided paired Wilcoxon signed-rank test (n=16 flies per pairwise test). Traces over time plotted as mean ± SEM.

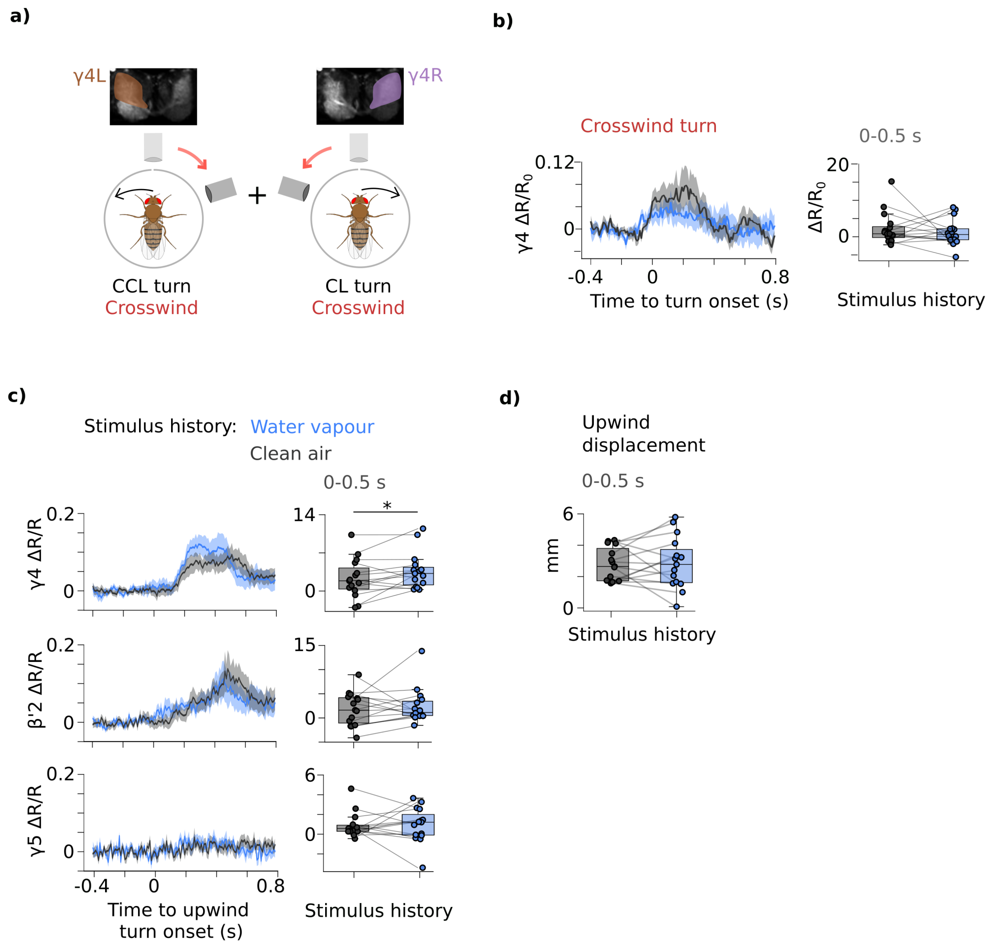


**Supplementary Figure 7. Stimulus history is not represented in bilateral β′2 or γ5 DAN activity. a,** For crosswind turns, γ4 DAN activity from the left hemisphere during counterclockwise (CCL) turns was combined with activity from the right hemisphere during clockwise (CL) turns, as done for upwind turns in Figure 6a. **b,** There was no effect of vapour history on crosswind turn-triggered ipsilateral γ4 DAN activity  (n=15 flies). c, There was no effect of vapour history on bilateral β′2 or γ5 DAN activity. Bilateral γ4 DAN activity reflects vapour history, but the increase. is less apparent as compared to combining ipsilateral activity as shown in Figure 6a-b (n=15 flies per paired group).  d) Upwind turn-triggered γ4 DAN activity increases following vapour exposure (Figure 6b, Supplementary Figure 7b), there is no difference in upwind approach (measured over 0-0.5 s following upwind turn onset; n=15 flies).  Statistical comparisons made using two-sided paired Wilcoxon signed-rank test.
